## Supplementary material for "Glial Nrf2 signaling mediates the neuroprotection exerted by *Gastrodia elata* Blume in Lrrk2-G2019S Parkinson’s disease": figure supplement

Figure Supplements for

*Correspondence:

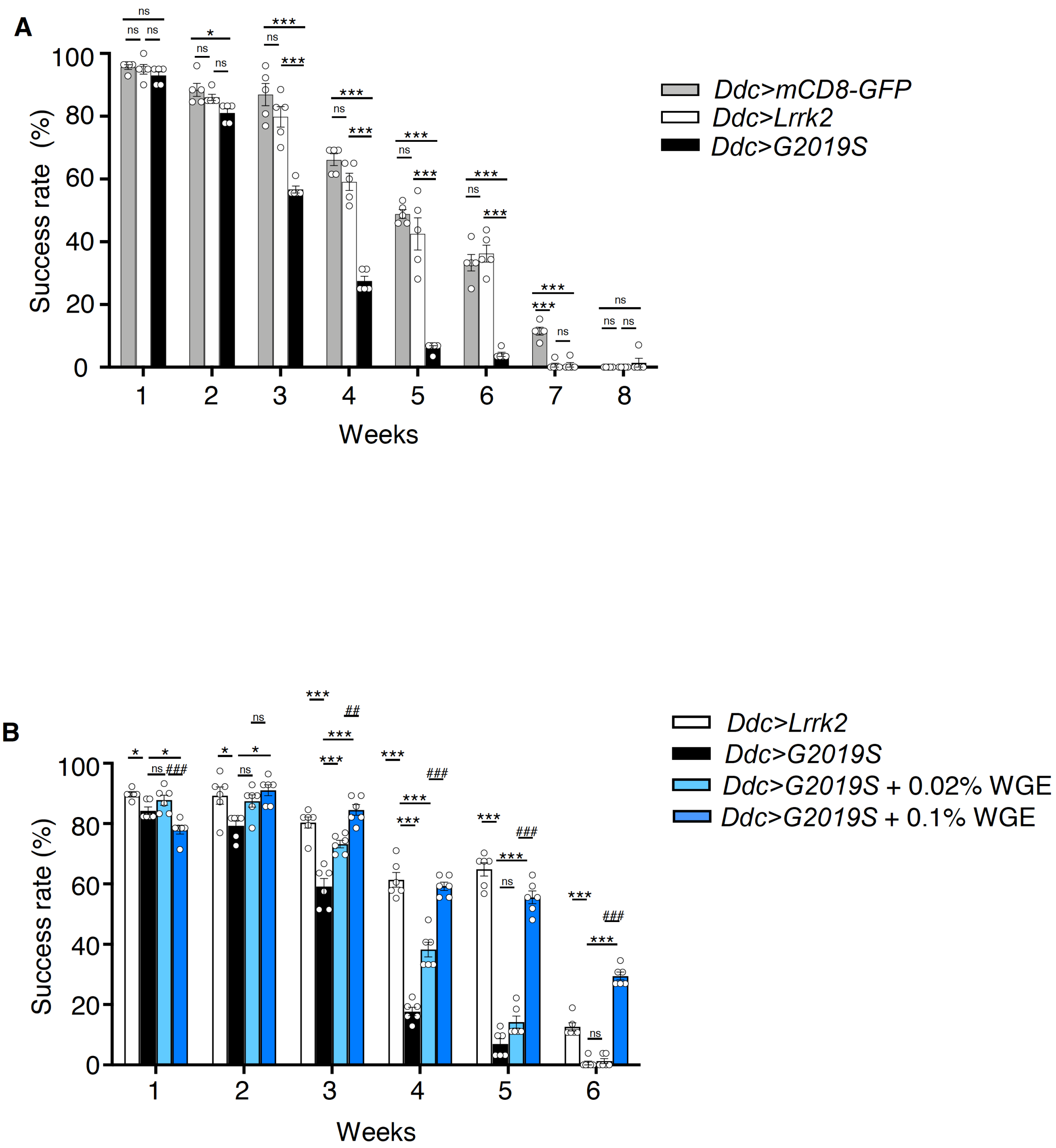


**Figure 1-figure supplement 1. Climbing activity assay of *Ddc>mCD8-GFP* and *Ddc>Lrrk2* flies.**

(**A**) Comparable climbing activities were detected for *Ddc>mCD8-GFP* and *Ddc>Lrrk2* flies. Both exhibited better climbing activities than *Ddc>G2019S* flies in the climbing assay from week 1 to 6. Bar graph shows percentage (mean ± SEM, N = 5) of flies that successfully climbed above 8 cm within 10 sec. One-way analysis of variance (ANOVA) and Tukey's post-hoc multiple comparison test: * $p < 0.05$, *** $p < 0.001$, ns, not significant. (**B**) *Ddc>G2019S* flies fed with 0.02% or 0.1% WGE exhibited improved climbing activity (mean ± SEM, N = 6). One-way ANOVA and Tukey's post-hoc multiple comparison test: * $p < 0.05$, *** $p < 0.001$ (relative to *Ddc>G2019S*); and ## $p < 0.01$, ### $p < 0.001$ (comparing different doses of WGE), ns, not significant.

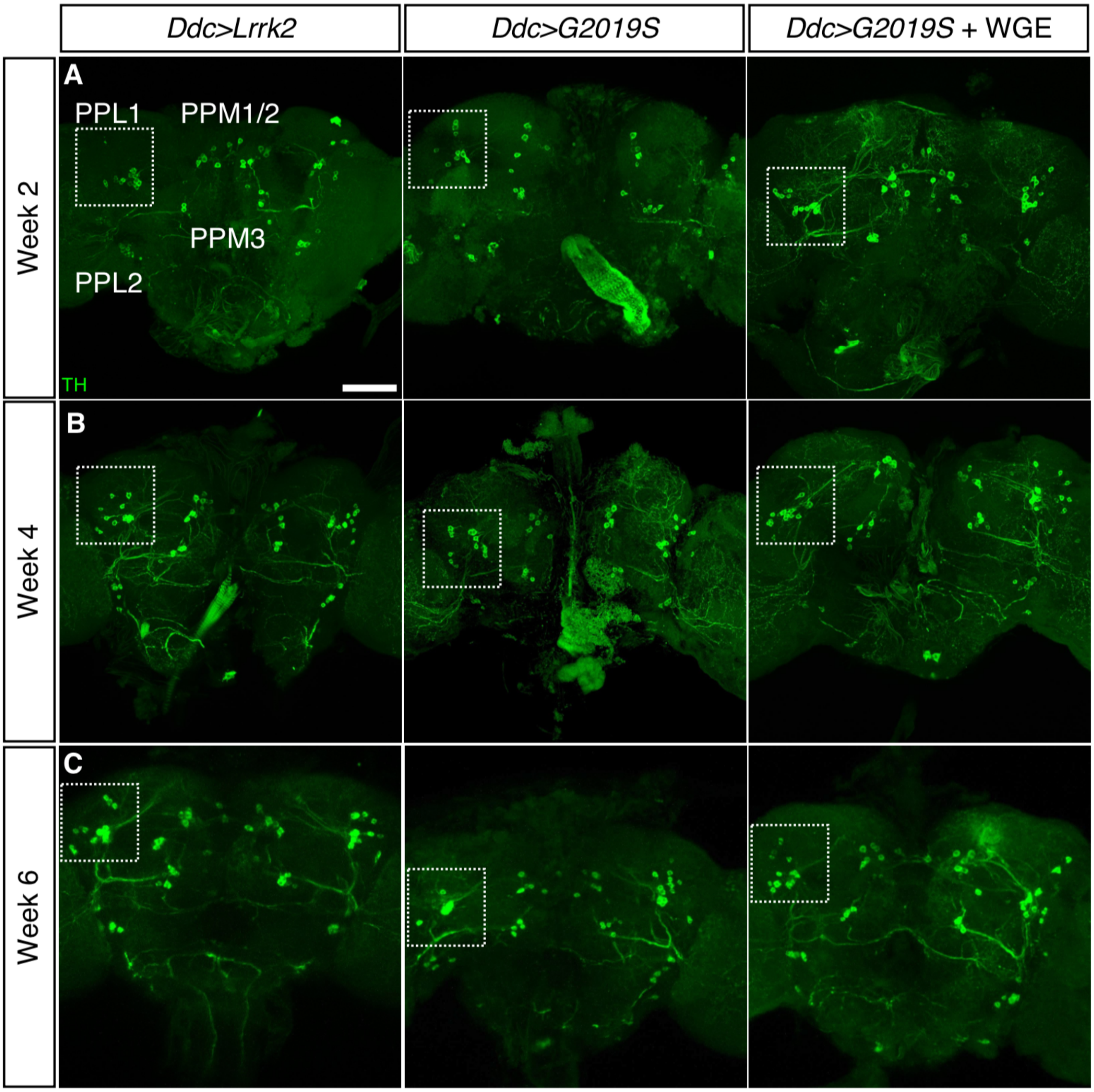


**Figure 2-figure supplement 1. WGE treatment prevents dopaminergic neuron loss in *Ddc>G2019S* flies.** (**A** to **C**) Representative adult whole-brain images for TH staining to reveal dopaminergic neurons in the PPL1, PPL2, PPM1/2 and PPM3 clusters of 2-, 4-, and 6-week-old flies. Scale bar: 40 μm. The images for the PPL1 cluster are shown as enhanced views of the dashed boxes in Fig. 2.

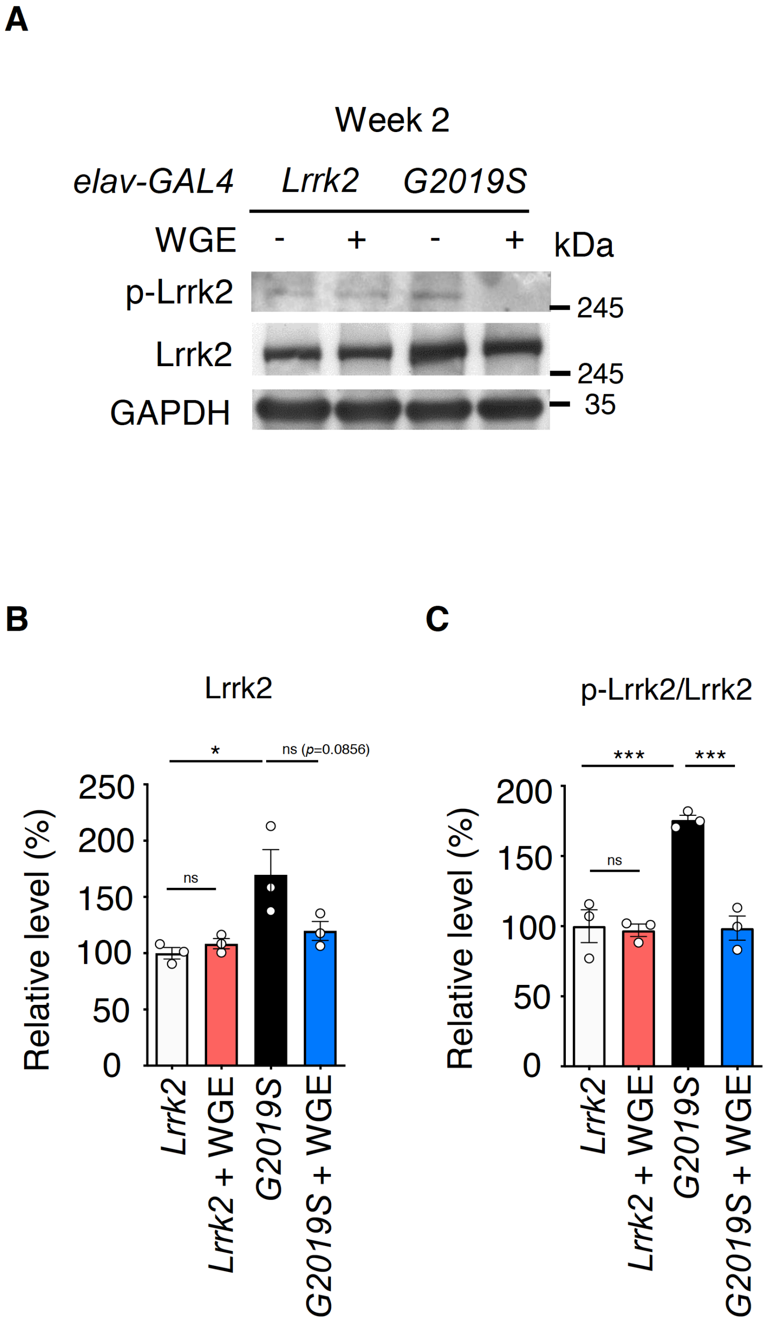


**Figure 3-figure supplement 1. WGE specifically modulates Lrrk2 accumulation and hyperactivation in *elav>G2019S* but not *elav>Lrrk2* flies.**

(**A**) Representative immunoblots of 2-week-old adult brain lysates showing levels of Lrrk2 and pLrrk2 ($Ser^{1292}$) in *elav>Lrrk2*, WGE-fed *elav>Lrrk2, elav>G2019S* and WGE-fed *elav>G2019S* flies. (**B** and **C**) Quantification (mean ± SEM, N = 3) of Lrrk2 (B) and pLrrk2/Lrrk2 (C) levels. One-way ANOVA and Tukey's post-hoc multiple comparison test: * $p < 0.05$, *** $p < 0.001$, ns, not significant.

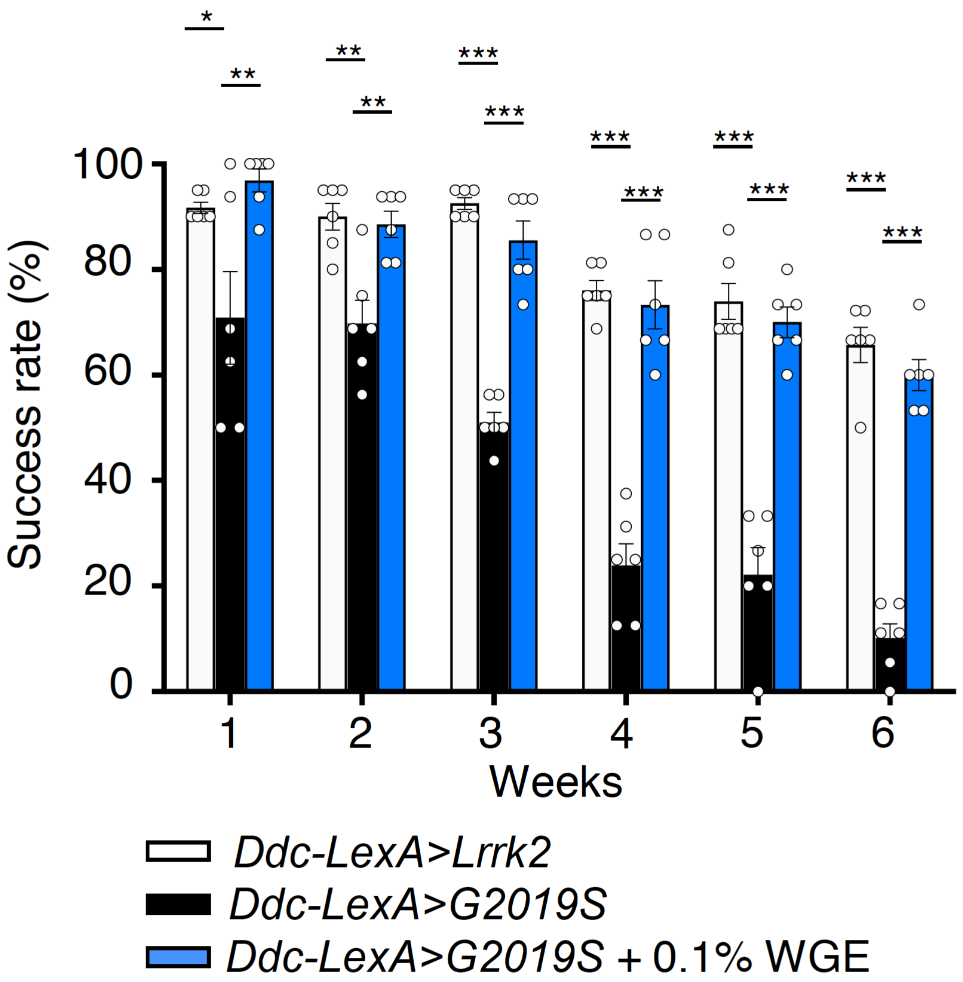


**Figure 5-figure supplement 1. WGE rescues the locomotion defect displayed by *Ddc-LexA>G2019S* flies.** WGE treatment improves the climbing ability of *Ddc-LexA>G2019S* flies. Bar graph shows percentage (mean ± SEM, N = 6) of flies that successfully climbed above 8 cm within 10 sec. One-way ANOVA and Tukey's post-hoc multiple comparison test (relative to *Ddc-LexA>G2019S*): * $p < 0.05$, ** $p < 0.01$, *** $p < 0.001$.

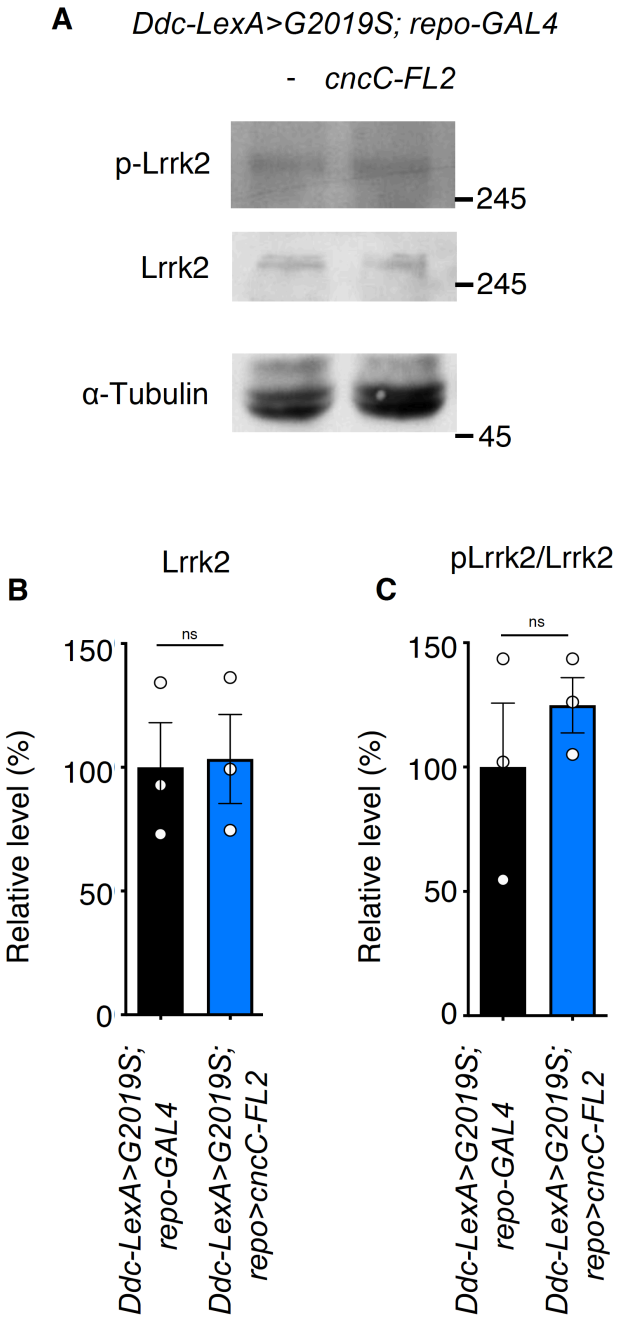


**Figure 6-figure supplement 1. Lrrk2 and pLrrk2 levels are maintained upon glial Nrf2 overexpression.** (**A**) Representative immunoblots of 2-week-old adult brain lysates showing protein expression levels of Lrrk2 and pLrrk2 ($Ser^{1292}$) in *Ddc-LexA>G2019S; repo-GAL4* and *Ddc-LexA>G2019S; repo>cncC-FL2* flies. (**B** and **C**) Quantification of Lrrk2 (B) and pLrrk2/Lrrk2 (C) (mean ± SEM, N = 3). Student *t*-test: ns, not significant.

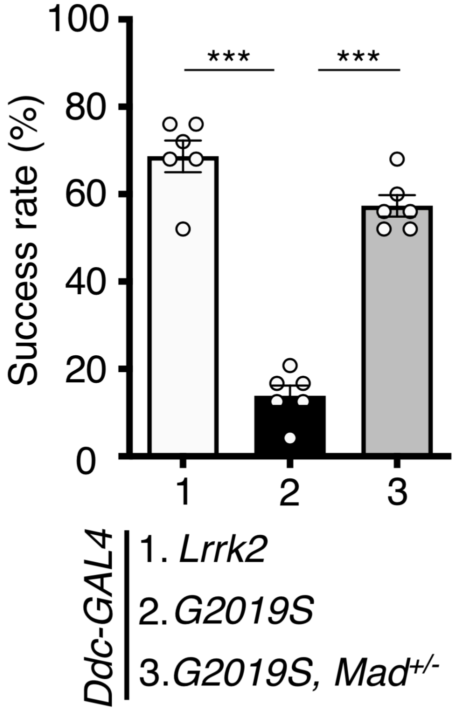


**Figure 7-figure supplement 1.** ***Mad*** **heterozygosity rescues the impaired locomotion of** ***Ddc>G2019S*** **flies.** Removing one copy of *Mad* improves *Ddc>G2019S* climbing activity. Bar graphs show climbing success rates (mean ± SEM, N = 6) of 6-week-old *Ddc>Lrrk2, Ddc>G2019S,* and *Ddc>G2019S, Mad*$^{-/+}$ flies. One-way ANOVA and Tukey's post-hoc multiple comparison test (relative to *Ddc>G2019S*): *** $p < 0.001$.
